# ColabDock: inverting AlphaFold structure prediction model for protein-protein docking with experimental restraints

**DOI:** 10.1101/2023.07.04.547599

**Authors:** Shihao Feng, Zhenyu Chen, Chengwei Zhang, Yuhao Xie, Sergey Ovchinnikov, Yiqin Gao, Sirui Liu

## Abstract

Prediction of protein complex structures and interfaces potentially has wide applications and can benefit the study of biological mechanisms involving protein-protein interactions. However, the surface prediction accuracy of traditional docking methods and AlphaFold-Multimer is limited. Here we present ColabDock, a framework that makes use of ColabDesign, but reimplements it for the purpose of restrained complex conformation prediction. With a generation-prediction architecture and trained ranking model, ColabDock outperforms HADDOCK and ClusPro not only in complex structure predictions with simulated residue and surface restraints, but also in those assisted by NMR chemical shift perturbation as well as covalent labeling. It further assists antibody-antigen interface prediction with emulated interface scan restraints, which could be obtained by experiments such as Deep Mutation Scan. ColabDock provides a general approach to integrate sparse interface restraints of different experimental forms and sources into one optimization framework.

## Introduction

Proteins are essential components in biological processes and mostly execute their functions through interactions with other molecules. Understanding the structure of protein complexes is fundamental in comprehending biological mechanisms, which provide valuable information for structure-based drug discovery. Several methods are available for determining the complex structure experimentally, including x-ray crystallography, nuclear magnetic resonance (NMR) spectroscopy, and cryo-electron microscopy (cryo-EM). Although widely used in the field of structural biology, each method has its own limitations and is costly and/or time-consuming to obtain structures, which necessitates efficient computational methods for predicting complex structures.

Protein-protein docking is one of the most important problems in computational biology. Given the individual structures of each component in the protein complex, a typical docking method generates a large number of complex conformations using a Fast Fourier Transform (FFT)-based algorithm, and evaluates the conformations with a customized scoring function. Various algorithms have been developed for this free docking method, such as ZDOCK [1], pyDock [2], SwarmDock [3], HADDOCK [4, 5], and ClusPro [6-8]. Despite that free docking methods have had some success, they suffer from the moderate accuracy of the scoring functions [9]. To alleviate this problem, restraints derived from experimental methods are incorporated in docking algorithms, since such methods are more accessible and easier to implement than structure determination methods. For example, chemical cross-linking (XL) provides the distance between two residues cross-linked by fixed-length reagents, whereas Nuclear Overhauser Effect (NOE) in NMR measures distances between atom pairs. Although these experimental restraints are sparse and cannot fully determine the protein complex structure, they may provide crucial information about the component interaction interface.

To incorporate the experimental data, many restrained docking algorithms have been developed, taking in experimental restraints with different strategies. HADDOCK [4, 5] allows users to define the active and passive residues in the protein complex. Active residues are those known to make contact within the complex, while passive residues only potentially make contact. The active and passive residues are then converted into the form of Ambiguous Interaction Restraints (AIRs), which serve as an energy term in optimization and conformation ranking. ZDOCK [1] utilizes contacting residues to filter docking conformations. pyDOCK [2] adopts a similar posteriori strategy, employing the percentage of satisfied restraints as a pseudo-energy term in the scoring function. ClusPro [6-8] generates a feasible translation set for each restraint and selects translations from the intersection set with frequency larger than a cutoff.

On the other hand, multiple deep neural network based models have been proposed for *ab initio* protein structure prediction, such as AlphaFold2 (AF2) [10], AlphaFold-Multimer [11], and RoseTTAFold2 [12]. It has been shown that AF2 has learned an approximate biophysical energy function from massive protein structure data and achieves state-of-the-art performance in protein model quality estimation [13]. Besides, AF2 has also been used in protein design by either Monte Carlo Markov Chain (MCMC) optimization [14-16] or by gradient backpropagation [17]. However, the accuracy of protein complex structure prediction models like AlphaFold-Multimer is limited in many cases, especially for flexible protein-protein interaction. The predicted structures from these models are not always consistent with the experimental observations.

Inspired by these studies, here we present ColabDock, a new protein-protein docking framework guided by sparse experimental restraints. ColabDock replaces the FFT algorithm with gradient backpropagation and automatically integrates the AF2 energy function and restraints posteriori. The framework contains two stages: the generation stage and the prediction stage. In the generation stage, ColabDesign [17], a protein design framework developed based on AF2, is adopted. The input sequence is optimized in the logit space to generate a complex structure that fits the individual structures of each component and the provided restraints, while maximizing pLDDT and pAE measures. In the prediction stage, the final structure is predicted using AF2 based on the generated complex structure and individual component structures. For each protein, ColabDock framework performs multiple runs and generates diverse conformations. The final conformation is selected by a ranking SVM algorithm. We evaluate ColabDock performance on a synthetic dataset and several types of experimental restraints, including NMR, covalent labeling, and simulated Deep Mutational Scanning (DMS). The results demonstrate that ColabDock achieves better performance than the FFT-based docking algorithms.

## Results

Due to the limited accuracy in complex structure prediction of deep learning based models such as AF2 and AlphaFold-Multimer, as well as the energy-based free docking methods, the predicted structures are not always consistent with the experimental restraints. Here, to solve the inconsistency, we propose ColabDock by incorporating the experimental restraints into protein complex structure modeling. ColabDock is a framework that fulfills accurate restrained protein interface prediction through optimizing the input sequence under the designed loss to generate complex structures that are in accord with the provided experimental restraints, and refining the overall structure with AlphaFold (Figure 1). This endows ColabDock with the flexibility to take various types of restraints as input. Here, we mainly focus on two types of restraints. The first type restrains the distance of a residue pair below a certain threshold. It is on the residue-residue level and is referred to as the 1v1 restraint hereafter. Restraints derived from Cross-Linking Mass Spectrometry (XL-MS) belong to this type. The second type provides a series of residues potentially located on the protein-protein interface for each chain, and it is possible that some of the detected interface residues do not interact with those from different chains due to chain multiplicity, surface coverage and noise. This type of restraint is on the interface level and is referred to as the MvN restraint. Examples include NMR and covalent labeling. To alleviate the influence of the stochasticity in initialization and optimization, we run ColabDock multiple times to generate diverse complex structures and use the proposed ranking algorithm to pick the best predictions. Experiments on ranking algorithm training and hyperparameter tuning on round and step numbers are performed on the development set with details described in support information (Figure S1). The default values of round for 1v1 and MvN restraints are set to 15 and 10, respectively. For the step parameter, it is set to 50. We use the default settings in all experiments described below. To demonstrate the effectiveness of ColabDock, we evaluate the performance of ColabDock on the validation and benchmark sets, and compare the performance to two state-of-the-art algorithms, i.e., HADDOCK and ClusPro, on the NMR Chemical Shift Perturbation (CSP), Covalent Labeling (CL), and simulated antibody-antigen sets.

**Figure 1.**
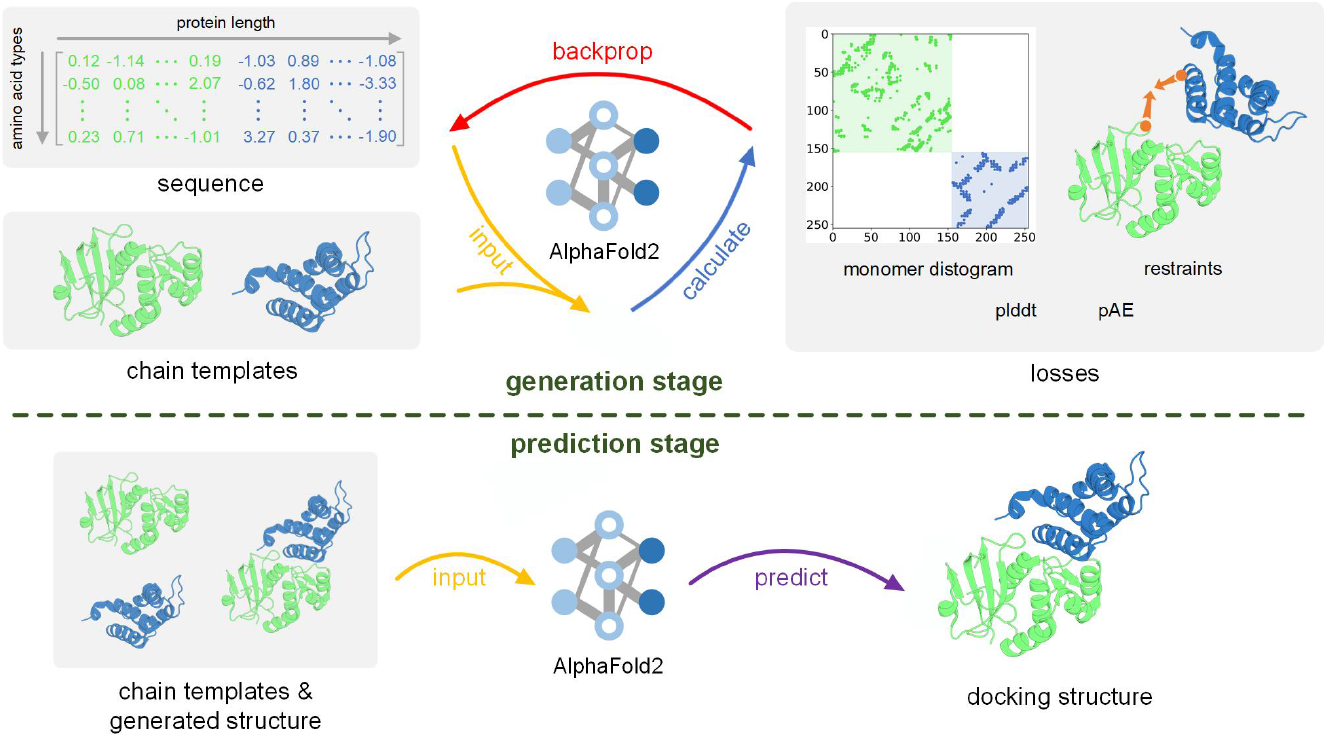
The workflow of ColabDock. ColabDock is a framework that incorporates experimental restraints into protein complex structure modeling. It consists of two stages. In the generation stage, ColabDock generates complex structures that are in accord with the provided experimental restraints and the input templates of each chain, through optimizing the input sequences in the logits space. In the prediction stage, the generated structure along with the input templates are fed into AF2 to make the final prediction.

### Validating the ColabDock performance with simulated 1v1 and MvN restraints

In this section, we first evaluate the performance of ColabDock on complexes with 1v1 and MvN restraints. Then, we compare the structures derived from the generation stage and the prediction stage, to evaluate the effect of the latter. DockQ [18] is a commonly used evaluation score for interface structure, and DockQ>0.23 indicates the interface quality is acceptable, regarded here as successful predictions for all methods evaluated (see Methods).

For each protein complex in the 1v1 restrained validation set, we randomly sample 2, 3, or 5 1v1 restraints to simulate the XL-MS experiment, and evaluate the maximal DockQs of the top1, top5 and top10 structures elected by the proposed ranking algorithm (see Methods) as well as the maximal DockQ of all predictions in a sample. Among the 111 samples, the maximal DockQs of 108 are above 0.23. For top1 and top10, the numbers are 101 and 105, respectively (Table 1). Although one might expect interfaces to be of lower quality with fewer restraints as less information is provided, under the most difficult circumstance where only 2 1v1 restraints are provided, the maximal DockQ of 81.08% protein complexes are above 0.23 (Figure 2a). For circumstances with 3 and 5 restraints, the success percentages are close to 100%, which demonstrates the framework’s ability to accurately predict and select high-quality structures. The ability to find the optimized conformation with only 2 restraints also indicates the ability of AF2 to search through the detailed interface potential as long as restraints can guide it to a near-correct conformation space.

**Table 1.** Performance of ColabDock on validation set.

| Restrains Type | Candidate Type | Mean DockQ | Mean RMSD(Å) | Success Rate<sup>a</sup> |
| --- | --- | --- | --- | --- |
| 1v1 Restrains | Top1<sup>b</sup> | 0.610 | 3.26 | 101/111 |
|  | Top5<sup>c</sup> | 0.660 | 2.87 | 104/111 |
|  | Top10<sup>d</sup> | 0.675 | 2.67 | 107/111 |
| MvN Restrains | Top1 | 0.443 | 5.53 | 75/111 |
|  | Top5 | 0.488 | 4.84 | 82/111 |
|  | Top10 | 0.505 | 4.55 | 83/111 |

a. Success rate is represented as the number of successful samples divided by total sample number. A sample prediction is regarded as successful if the DockQ of the best candidate structure is greater than 0.23.
b. Top1 refers to the first structure ranked according to the docking method.
c. Top5 refers to the top five structures ranked according to the docking method.
d. Top10 refers to the top ten structures ranked according to the docking method.

**Figure 2.**
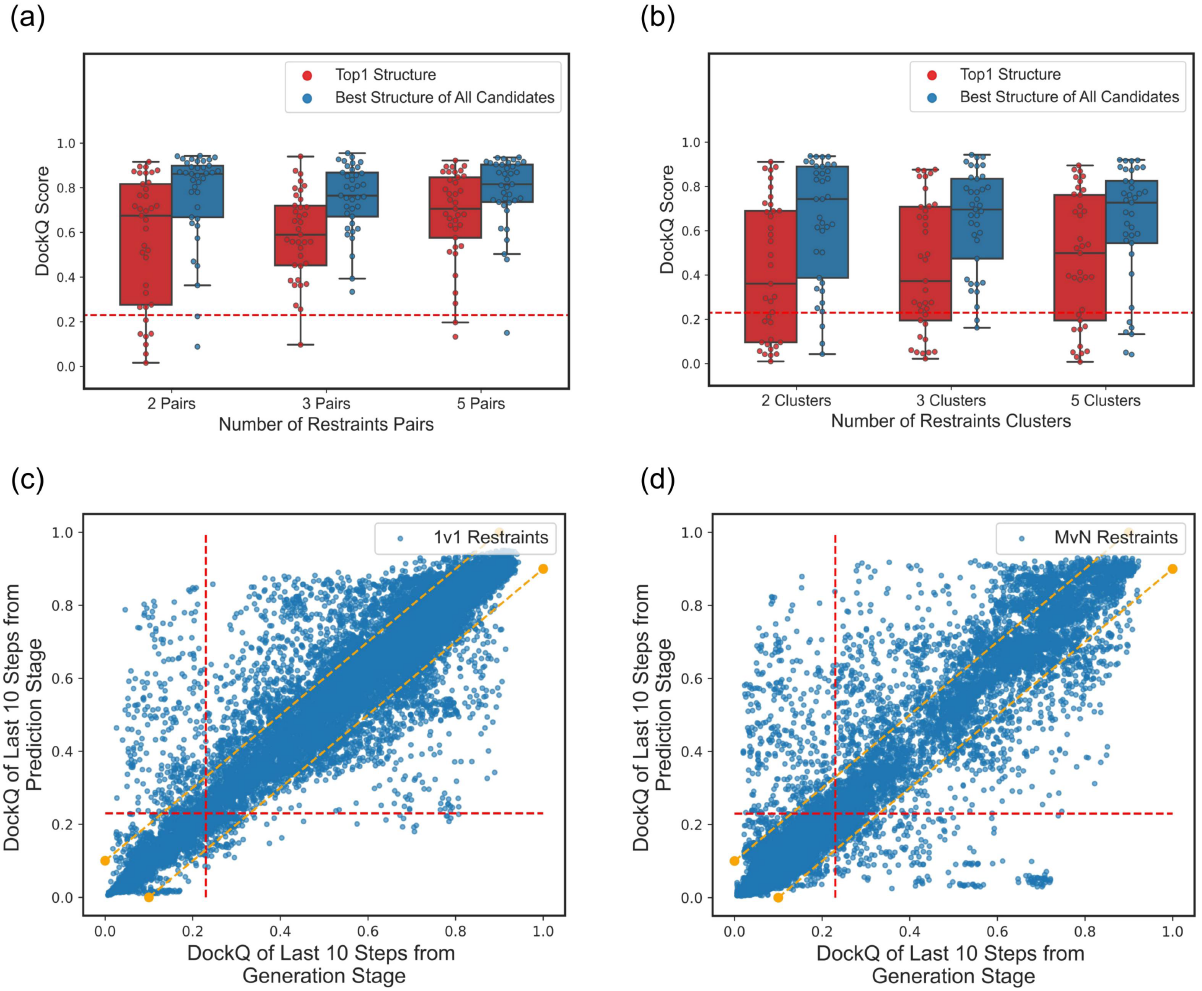
Performance of ColabDock on the validation set. **a**. The DockQ distribution of ColabDock predictions with 1v1 restraints. **b**. The DockQ distribution of ColabDock predictions with MvN restraints. **c**. and **d**. are DockQ comparisons on structures derived from the generation and prediction stage in ColabDock, with 1v1 restraints (**c**) and MvN restraints (**d**). Data points above the upper yellow line means the DockQ of the prediction stage is at least 0.1 larger than that of the generation stage, and vice versa.

To further assess the model performance on the MvN restraints, we evaluate it on the same 111 proteins with the MvN restraints converted from the 1v1 ones (see Methods). This task emulates the NMR Chemical Shift Perturbation or Paramagnetic Relaxation Enhancements (PRE) as well as covalent labeling data, and is more challenging since it provides surface instead of residue pair level correspondence. As are shown in Table 1 and Figure 2b, out of 111 complexes, the maximal DockQs of 100 complexes are above 0.23. For top1 structures, the number is equal to 75. The average DockQ and RMSD of top1 structures are 0.443 and 5.534, respectively. These results demonstrate ColabDock can also deal with vague restraints and achieve satisfying performance. For different restraint numbers, as restraints’ quality increases, the model is able to pick out more correct structures (Figure 2b) and the success rate for top1 ranked prediction (defined as the correct structure number of top1 candidates divided by the number of subsamples) increases from 62.16% (2 pairs) to 70.27% (3 pairs) and 70.27% (5 pairs).

To evaluate the necessity of the prediction stage in ColabDock, based on the complex structures generated in the two experiments on 1v1 and MvN restraints, we compare the performance of the generation stage and the prediction stage. Specifically, we collect the structures from the last 10 steps of each round, by which most optimization has reached convergence. For protein complexes with 1v1 restraints, the mean DockQ of the structures derived from the generation stage is 0.554, lower than that of the prediction stage (0.572). For MvN restraints, the performance is similar. The mean DockQs of the generation stage and the prediction stage are 0.266 and 0.276, respectively. For cases in which the interface difference in generation and prediction stages is sufficiently large, defined as DockQ differences larger than 0.1 here, the prediction stage archives better performance on 69.9% complexes with 1v1 restraints (Figure 2c). On MvN restraints, the percentage is 68.8% (Figure 2d). These results suggest the learned potential by AlphaFold in the prediction stage can help fix the complex structure derived from the generation stage and justifies the presence of the prediction stage.

### ColabDock outperforms traditional docking methods on the benchmark set

We construct a protein-protein benchmark set from protein docking benchmark 5.5 (see Methods for details). It contains 37 common protein complexes on which the DockQs of the structures predicted by AlphaFold-Multimer are below 0.23. This dataset is complementary to the validation and development sets used above. We compare ColabDock with two representative docking methods, HADDOCK and ClusPro. Both of the docking methods performed excellently in the CAPRI (Critical Assessment of PRediction of Interactions). HADDOCK is specially designed to integrate experimental restraints for protein complexes modeling. We followed default settings for HADDOCK and ClusPro with minor adjustment (see detailed settings in SI). For targets with lengths above 700 residues, we use segment-based optimization in ColabDock to avoid out of memory issues.

For each protein complex in the benchmark set, we sample 2,3, and 5 pairs of 1v1 restraints to guide docking, resulting in 111 benchmarking samples. ColabDock outperforms HADDOCK and ClusPro on most of the samples in the protein-protein benchmark set (Table 2, Figure 3a and Figure S2a). The mean DockQ of ColabDock is 0.477, and those of HADDOCK and ClusPro are 0.287 and 0.190, respectively. The mean RMSD of ColabDock is 5.00Å, also better than those of HADDOCK (7.16Å) and ClusPro (8.99Å). For different numbers of 1v1 restraints, ColabDock always achieves the best performance among the three methods (Figure 3b). Even with only 2 restraints, 23 out of 37 ColabDock predictions reach at least acceptable quality, while the numbers for HADDOCK and ClusPro are 14 and 7, respectively. These results suggest that ColabDock has the potential to generate native-like structures with even very sparse restraints, consistent with the observations on the validation set.

**Table 2.** Performance of three docking methods on synthetic protein-protein benchmark set.

| Restraints Type | Method | Mean DockQ | Mean RMSD(Å) | Success Rate<sup>a,b</sup> |
| --- | --- | --- | --- | --- |
| 1v1 Restraints | ColabDock | **0.477** | **5.00** | **88/111** |
|  | HADDOCK | 0.287 | 7.16 | 53/111 |
|  | ClusPro | 0.191 | 8.99 | 40/111 |
| MvN Restraints | ColabDock | **0.235** | **8.97** | **35/104** |
|  | HADDOCK | 0.132 | 10.45 | 17/104 |
|  | ClusPro | 0.114 | 11.37 | 13/104 |
| 1v1 Loose Restraints | ColabDock | **0.344** | **6.55** | **103/176** |
|  | HADDOCK | 0.234 | 6.83 | 75/176 |
|  | ClusPro | 0.174 | 8.49 | 51/176 |

a. Success rate is represented as the number of successful samples divided by total sample number. A sample prediction is regarded as successful if the DockQ of the best candidate structure is greater than 0.23.
b. We exclude 7 samples with MvN restraints and 9 samples with 1v1 loose restraints, on which ClusPro fails to give a prediction.

**Figure 3.**
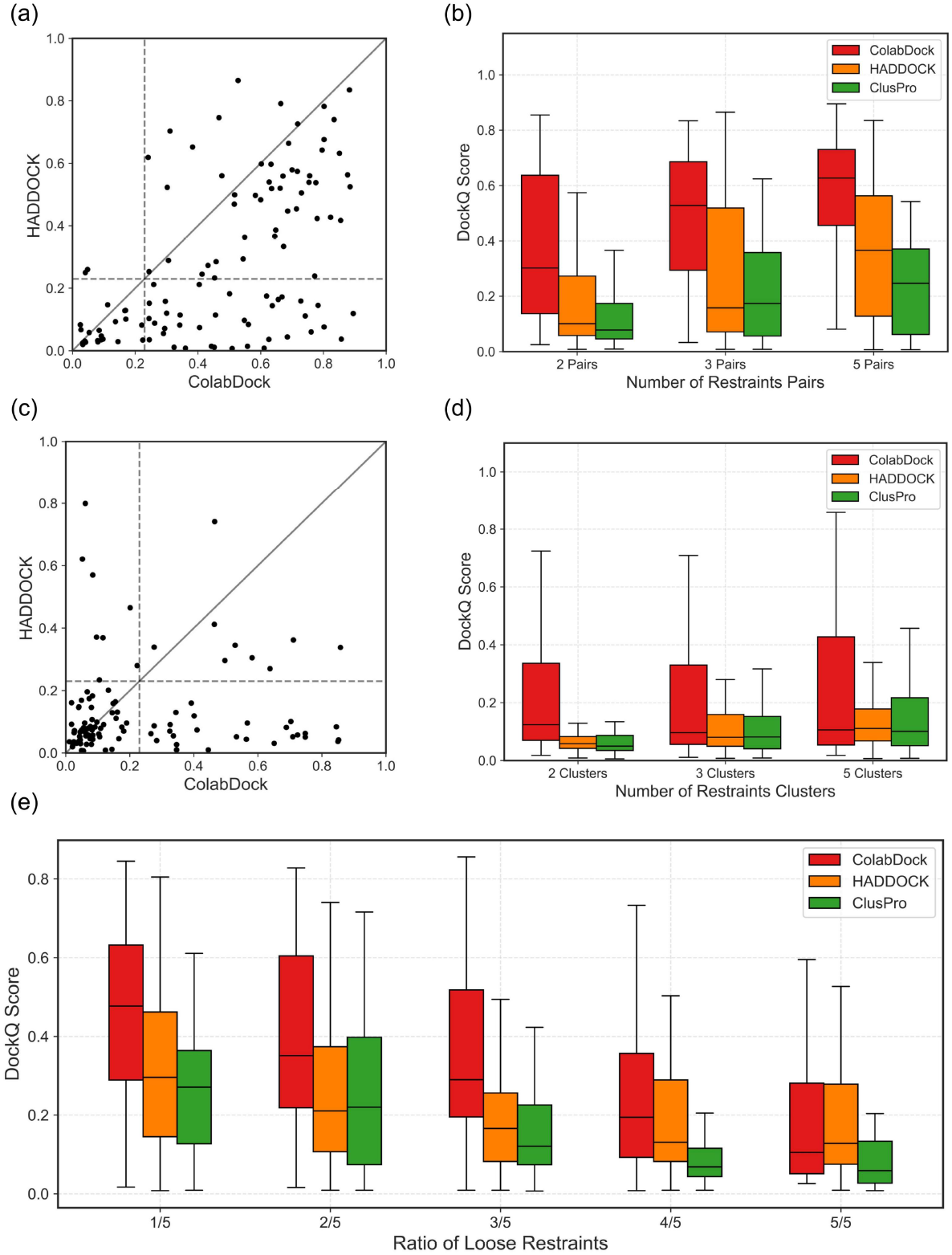
Comparison of ColabDock, HADDOCK, and ClusPro on the protein-protein benchmark set. **a**. DockQ scatter plot of top1 structures predicted by ColabDock and HADDOCK with 1v1 restraints. Two dotted lines are x=0.23 and y=0.23, respectively. Most of the structures predicted by ColabDock have a higher DockQ than those predicted by HADDOCK. **b**. The DockQ distributions of three docking methods with different numbers of 1v1 restraints. ColabDock beats the other two docking methods on 2, 3, and 5 1v1 restraints. **c**. DockQ scatter plot of top1 structures predicted by ColabDock and HADDOCK, with MvN restraints. ColabDock successfully predicts more samples (DockQ>=0.23) than HADDOCK. **d**. The DockQ distribution of three docking methods with different numbers of MvN restraints clusters. Increasing the number of restraints clusters can improve the modeling accuracy of ColabDock to some extent. **e**. The DockQ distribution of three docking methods with the ratio of loose restraints increasing. ColabDock holds the best performance in terms of median DockQ even with 4 out of 5 loose restraints.

Similar to the experiments on the validation set, we convert the 1v1 restraints above to MvN restraints to further evaluate the performance on interface level restraints. Since ClusPro fails to give predictions on 7 out of the 111 samples, we exclude corresponding samples from all three methods and make comparisons on the remaining 104 samples. Compared to the performance on complexes with 1v1 restraints, the performances of ColabDock, HADDOCK and ClusPro on MvN restraints decline at different degrees (Table 2), as a result of the ambiguity of the MvN restraints. Under this condition, ColabDock still outperforms the other two methods on DockQ (Table2, Figure 3c, and Figure S2b). The mean DockQ of ColabDock is 0.235, higher than those of HADDOCK and ClusPro, which are 0.132 and 0.114, respectively. In terms of mean RMSD, ColabDock achieves 8.97Å while HADDOCK and ClusPro are 10.45Å and 11.37Å. Among the 104 samples, ColabDock predicts acceptable structures for 35 samples, which is 2-3 times higher than that of HADDOCK (17) and ClusPro (13). Moreover, ColabDock always achieves the best performance on the DockQ metric no matter how much the cluster number of the MvN restraints is (Figure 3d).

It is common for restraints derived from experiments to include residue pairs far from the contact interface. To test the tolerance of ColabDock for looser restraints, we intentionally sample restraints at the distance range of 8-20Å, with the loose restraint number varying from 1 to 5 while the total restraint number is fixed at 5. We use these sampled restraints to guide the docking. ClusPro fails on 9 cases with 1v1 loose restraints and we exclude corresponding samples of all three methods to evaluate the performance. As summarized in Table 2, ColabDock shows the best performance on the 176 samples integrated with 1v1 loose restraints where the mean DockQ is 0.344 and the mean Cα-RMSD is 6.55Å (Figure S2c-d, Figure 3e). Even when 3 out of 5 restraints are loose restraints, ColabDock still reaches a mean DockQ score of 0.363 while the median DockQs of HADDOCK and ClusPro reduce to 0.211 and 0.172, respectively. Meanwhile, with the number of looser restraints decreasing, the performance of ColabDock shows a significant improvement compared to that of HADDOCK and ClusPro. These results indicate that the performance of ColabDock relies weakly on the quality of restraints. But when integrated with high quality distance restraints, ColabDock can generate complex structures closer to the ground truth than the other two methods.

Above results indicate that ColabDock is a robust and highly accurate method to predict protein complex structure and can utilize different types of restraint information effectively. This method can serve as an alternative approach to generate native-like protein complex structures that utilizes information from restraints and fits well with the provided experimental evidence. Moreover, with the average number of surface residues increasing, ColabDock is able to improve the interface to higher proportion and quality.

### ColabDock improves interface and overall structure prediction on the NMR derived CSP restraints

Chemical Shift perturbation (CSP) is an NMR surface detection experiment. It cannot specifically identify which two residues are interacting, but instead provides a range of residues located at the interface. This kind of restraint information therefore falls into the category of MvN restraints. We obtain two experimental protein complex samples with CSP information from Dominguez [5] (see CSP set in the Methods), and compare the performance of ColabDock, HADDOCK, and ClusPro on them.

In both cases, the interface structures generated by the ColabDock model exhibit high quality. The DockQ scores reach 0.603 and 0.936, respectively, significantly higher than those of HADDOCK (EIN: 0.385, E2A: 0.411) and ClusPro (EIN: 0.347, E2A: 0.131) (Table 3). The top5 and top10 ranking structures from ColabDock still outperform those of HADDOCK and ClusPro in both average DockQ and Cα-RMSD (Table 3 and Figure S3a). Meanwhile, the low variance of the top10 structures predicted by ColabDock indicates the consistency with the experimental restraints (Figure 4a).

**Table 3.**
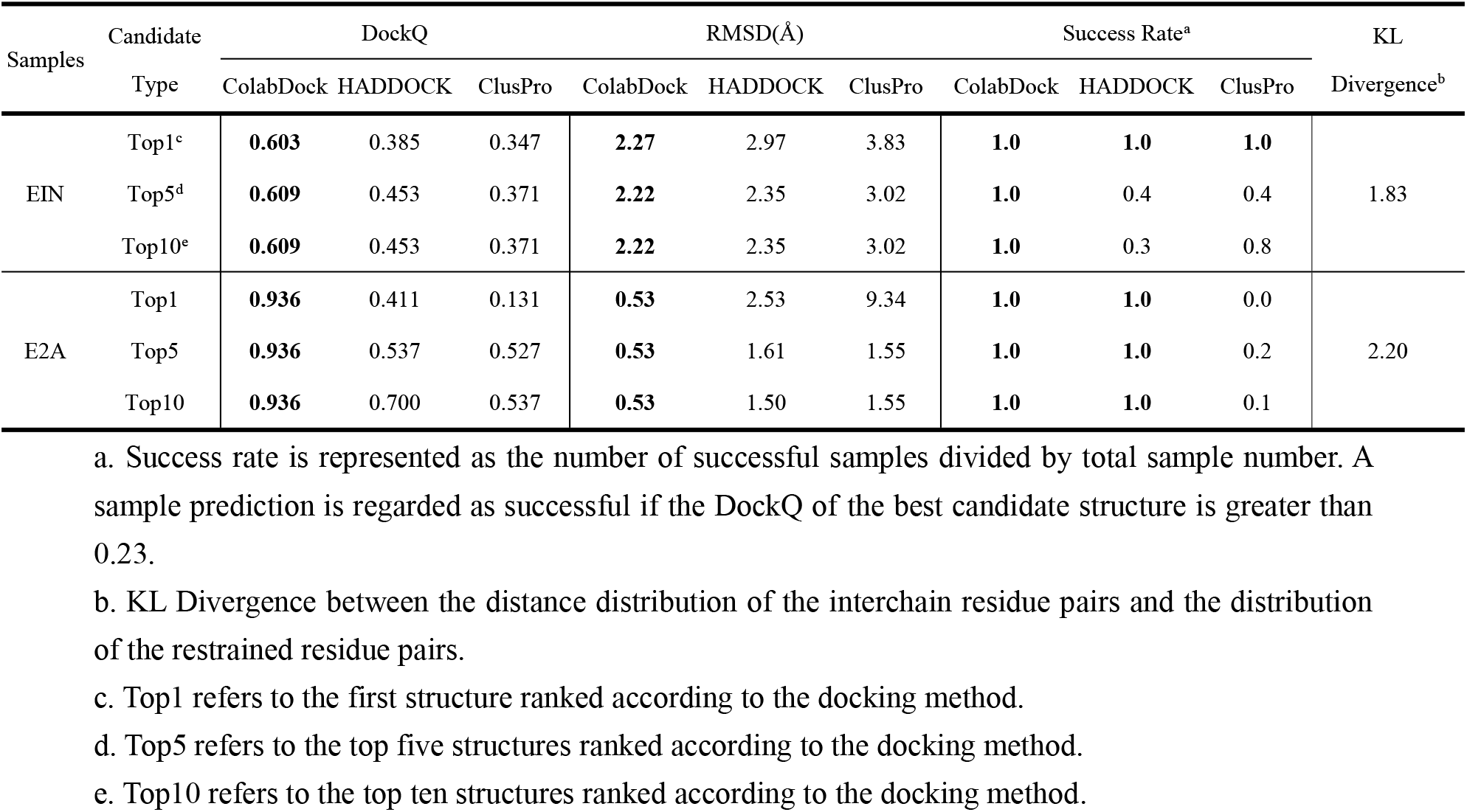
Performance of three docking methods on NMR CSP restraints.

**Figure 4.**
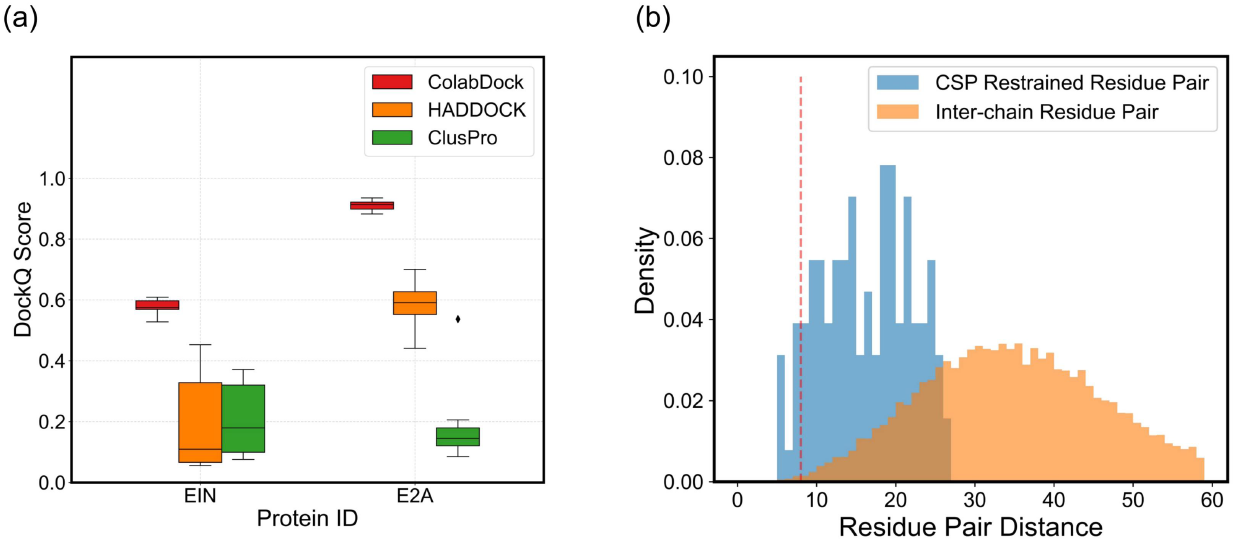
ColabDock performance and restraints analysis on NMR CSP restraints. **a**. DockQ distributions of top10 structures predicted by ColabDock (red), HADDOCK (orange), and ClusPro (green) on the two complexes with CSP restraints. **b**. The distance distributions of interchain residue pairs (orange) and the experimental restraints (blue) in protein EIN. The KL divergence of the two distributions is 1.83.

We also evaluated the quality of the CSP restraints. We calculated the Kullback-Leibler (KL) divergence between the distance distribution of all residue pairs given by experiment and that of all inter-chain residue pairs, to compare the two distributions. A higher KL divergence indicates two distributions are different from each other, which implies the restraints provide more interface information. KL divergences of the two samples are 1.83 and 2.20 (Table 3), and their corresponding distributions are illustrated in Figure 4b and Figure S4a, respectively. These statistics demonstrate the high quality of NMR restraints. The performance of ColabDock suggests its higher ability to handle high quality MvN restraints than the other two methods.

### Performance of ColabDock on varied CL restraints

Covalent labeling (CL) can be used to label the side chain of residue with reagents. Residues with a significant modification ratio change between bounded and unbounded states are more likely to locate in the interacting surface in the protein complex. CL restraints can be classified as MvN restraints. Typically, the CL restraints possess a wider distance range than CSP restraints and provide relatively less interface information.

We benchmark ColabDock on 5 protein assemblies with covalent labeling restraints available from Drake [19] (see CL set in the Methods), along with HADDOCK and ClusPro. ColabDock surpasses the other two methods on most of the assemblies (Table 4). On the top1 structure, the mean Cα-RMSD of ColabDock is 2.56Å, while those of HADDOCK and ClusPro are 5.04Å and 13.67Å, respectively. Additionally, ColabDock achieves the highest number of correct predictions (DockQ>=0.23) across all the top rankings (Figure 5a).

**Table 4.** Performance of three docking methods on covalent labeling restraints^a^.

| Samples | Candidate Type | DockQ |  |  | RMSD(Å) |  |  | Success rate |  |  | KL Divergence |
| --- | --- | --- | --- | --- | --- | --- | --- | --- | --- | --- | --- |
|  |  | ColabDock | HADDOCK | ClusPro | ColabDock | HADDOCK | ClusPro | ColabDock | HADDOCK | ClusPro |  |
| 4INS4 | Top1 | 0.627 | **0.734** | 0.033 | 2.32 | **1.74** | 10.69 | **1.0** | **1.0** | 0.0 | 0.32 |
|  | Top5 | 0.652 | **0.734** | 0.185 | 2.24 | **1.74** | 6.87 | **1.0** | 0.4 | 0.0 |  |
|  | Top10 | 0.659 | **0.752** | 0.304 | 2.20 | **1.69** | 3.74 | **1.0** | 0.5 | 0.1 |  |
| 4INS8 | Top1 | **0.332** | 0.176 | 0.027 | **4.45** | 6.51 | 15.11 | **1.0** | 0.0 | 0.0 | 0.16 |
|  | Top5 | **0.342** | 0.176 | 0.060 | **4.29** | 6.51 | 8.68 | **1.0** | 0.0 | 0.0 |  |
|  | Top10 | **0.373** | 0.176 | 0.133 | **4.13** | 6.51 | 7.56 | **0.8** | 0.0 | 0.0 |  |
| 4INS12 | Top1 | **0.812** | 0.247 | 0.040 | **1.27** | 4.92 | 11.51 | **1.0** | **1.0** | 0.0 | 0.16 |
|  | Top5 | **0.920** | 0.748 | 0.082 | **0.76** | 1.59 | 10.67 | **0.8** | 0.6 | 0.0 |  |
|  | Top10 | **0.920** | 0.748 | 0.082 | **0.76** | 1.58 | 8.91 | **0.8** | 0.6 | 0.0 |  |
| 2F8O | Top1 | **0.300** | 0.044 | 0.014 | **3.30** | 8.58 | 17.24 | **1.0** | 0.0 | 0.0 | 2.30 |
|  | Top5 | **0.332** | 0.091 | 0.059 | **3.11** | 8.58 | 9.58 | **0.6** | 0.0 | 0.0 |  |
|  | Top10 | **0.332** | 0.091 | 0.091 | **3.11** | 8.58 | 9.58 | **0.3** | 0.0 | 0.0 |  |
| 1YAG | Top1 | **0.710** | 0.318 | 0.035 | **1.47** | 3.43 | 13.82 | **1.0** | **1.0** | 0.0 | 0.56 |
|  | Top5 | **0.736** | 0.318 | 0.769 | **1.40** | 3.43 | 1.73 | **0.4** | 0.2 | 0.2 |  |
|  | Top10 | **0.736** | 0.318 | 0.769 | **1.40** | 3.43 | 1.73 | **0.2** | 0.1 | 0.1 |  |

a. The meanings of success rate, KL Divergence, top1, top5, and top10 are the same as Table 3.

**Figure 5.**
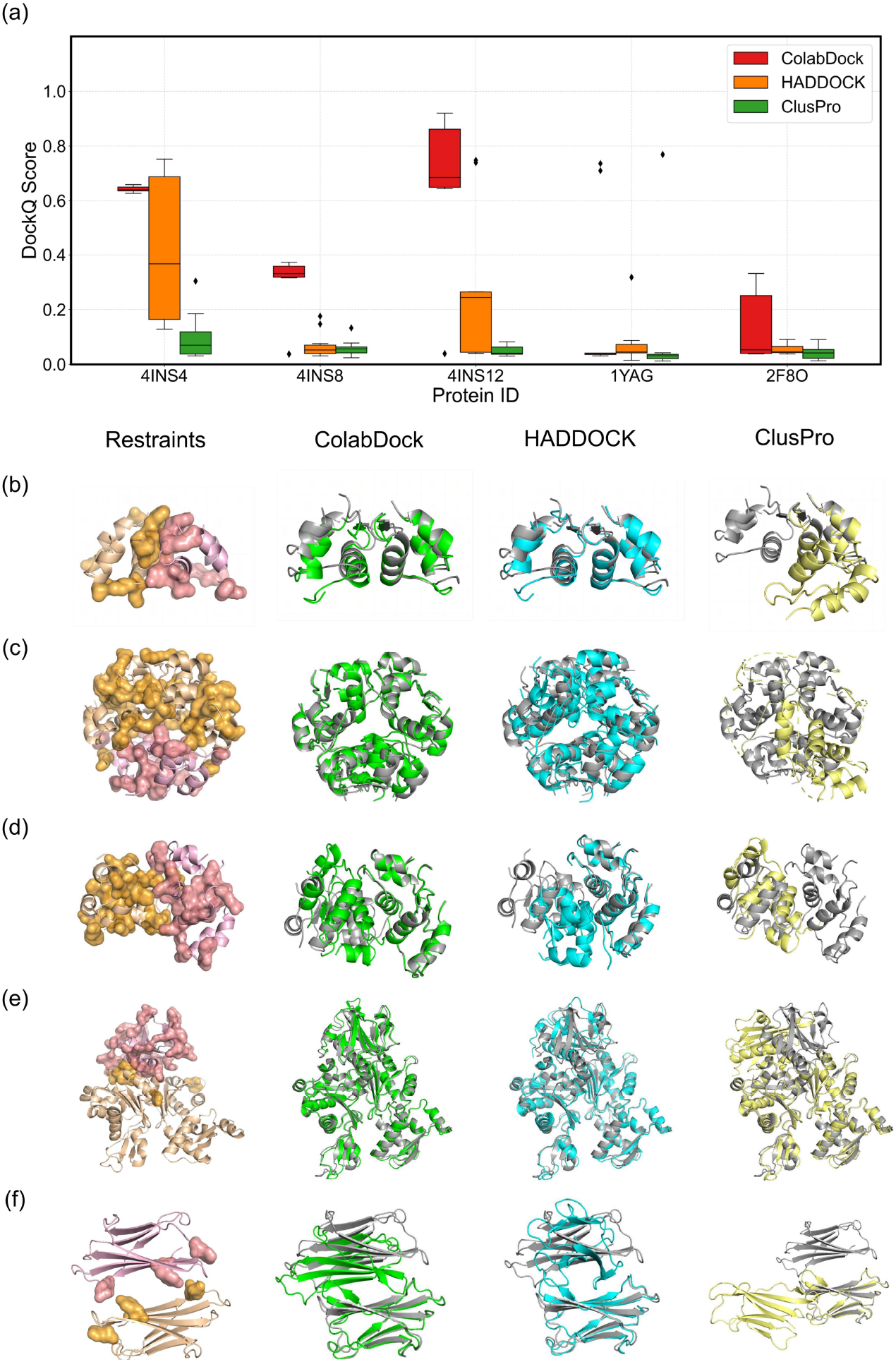
ColabDock performance and restraints analysis on covalent labeling samples. **a**. DockQ distributions for the top10 structures predicted by ColabDock (red), HADDOCK (orange), and ClusPro (green) on the five complexes with experimental CL restraints. **b**. 3D crystal structure of protein complex 4INS4, along with the predicted structures. From left to right are the crystal structure (two chains are colored in orange and pink, and CL restraints are colored in bright orange and salmon, respectively), the top1 structures predicted by ColabDock (green), HADDOCK (cyan), and ClusPro (yellow). Crystal structure is colored in gray in the latter 3 subfigures. 3D structures of crystal and predicted **c**. 4INS12, **d**.4INS8, **e**.2F8O, and **f**.1YAG follow the same color and order as 4INS4.

We next evaluate the quality of the restraints as in CSP restraints. It shows that the KL-divergence is lower in CL restraints, which indicates the less interface information contained in the restraints. We analyze the performance for each sample in detail. In circumstances where CL can provide high-quality interface information, such as in 4INS4 (Figure 5b, and Figure S4b) or 4INS12 (Figure 5c and Figure S4c), both ColabDock and HADDOCK can generate high-quality structures. For instance, in the top10 structures of 4INS4 predicted by ColabDock, the variances of the DockQs and RMSDs are less than those by HADDOCK (9.30×10^−5^ vs. 7.8×10^−2^ on DockQ and 1.16×10^−3^Å vs. 7.04Å on RMSD) (Figure 5a, Figure S3b). ColabDock achieves a higher average DockQ (ColabDock 0.642, HADDOCK 0.420, ClusPro 0.100) and a lower average RMSD (ColabDock 2.256Å, HADDOCK 4.552Å, ClusPro 9.388Å). This observation is consistent with that in the CSP dataset.

On protein 4INS8, although the experimental restraints are not primarily distributed on the interface (Table 4, Figure 5d), ColabDock still generates acceptable structures (Figure S4d), while HADDOCK and ClusPro produce incorrect conformations. When the distances of the experimental restraints are larger than 10Å, such as protein 2F8O (Figure 5e and Figure S4e), ColabDock can still predict acceptable structures, outperforming HADDOCK and ClusPro. The DockQs of the top1 predicted structure of ColabDock, HADDOCK and ClusPro are 0.30, 0.044, and 0.014, respectively.

In protein 1YAG, seven residues in chain A are known to be in contact with chain B, according to the experimental restraints. But no more information is provided about the contacting residues in chain B (Figure 5f, colored bright orange, Figure S4f). To deal with this, residues with solvent accessibility >= 40% are selected from chain B using FreeSASA software [20]. These residues serve as the contacting residues in chain B (Figure 5f, colored salmon). We then evaluate the performance of ColabDock, HADDOCK and ClusPro on this case. As are shown in Figure 5a and 5f, ColabDock and HADDOCK are both able to produce protein structures of satisfying quality, with the predictions of ColabDock in better agreement with the experimental restraints than HADDOCK. The DockQ of the top1 ranking structure predicted by ColabDock is 0.710, while that of HADDOCK is 0.318. As for ClusPro, although a structure of DockQ 0.769 is generated during the computation, ClusPro fails to select it as the top1 structure (Table 4). Additionally, the RMSD of the top1 structure of ColabDock (1.47Å) is lower than that of the best structure predicted by ClusPro (1.73Å).

For the five covalent labeling examples, despite the differences in the quantity and quality of the experimental restraints, ColabDock outperforms the two compared methods in most of the cases. Such a result suggests that ColabDock can effectively utilize the crucial information provided by experimental restraints and generate structures of high quality.

### Structure prediction for antibody-antigen complexes

Antibody-antigen complex modeling has been a long-standing challenge because of the flexibility of the CDR loop region and the lack of coevolution signal. In the protein docking benchmark 5.5, among the 67 antibody-antigen complexes, the DockQs of the predicted structures of 45 complexes are below 0.23. Here, we evaluate the performance on these failed complexes to find whether ColabDock can improve the model accuracy. Immunologists often use experiments like Deep Mutation Scanning (DMS) to determine which residues are more likely to participate in the antibody-antigen binding. Therefore, we simulate restraints derived from DMS by sampling residues on the interface (see Methods). For each complex in the antibody-antigen benchmark set, we sample 3 sets of different surface restraints. We use the sampled restraints to guide the docking of the 45 complexes and again evaluate the performance of ColabDock, HADDOCK, and ClusPro. Since most of the interaction surface residues on the antibody side are in the VH and VL region, we crop the antibody to only preserve the VH and VL region. For those with lengths longer than 1200, we also trim the antigen sequence to domains which are in direct contact with the antibody. All the three docking methods use the same sequences and restraints for prediction. ClusPro fails in four cases and for fairness we exclude the corresponding cases for all three methods in evaluation.

The results are summarized in Table 4. ColabDock is better than HADDOCK and ClusPro on the antibody-antigen benchmark, with a mean DockQ score of 0.223 and RMSD of 9.97 (Table 5, Figure 6a, Figure S5a-b). 43 out of 131 ColabDock top1 predictions reach at least acceptable quality while only 32 HADDOCK top1 predictions and 28 ClusPro top1 predictions meet this criterion. The number of at least medium quality cases (DockQ>=0.49) of ColabDock is apparently higher than that of HADDOCK and ClusPro (Figure 6b). Almost half (21) of the 43 successful ColabDock samples are of medium quality and above. In contrast, only 11 HADDOCK top1 results and 6 ClusPro are of at least medium quality.

**Table 5.** Performance of three docking methods on synthetic antibody-antigen benchmark set.

| Restrains Type | Method | Mean DockQ | Mean RMSD(Å) | Success Rate<sup>a,b</sup> |
| --- | --- | --- | --- | --- |
| Surface Restrains | ColabDock | **0.223** | **9.57** | **43/131** |
|  | HADDOCK | 0.181 | 9.97 | 32/131 |
|  | ClusPro | 0.136 | 11.37 | 28/131 |

a. Success rate is represented as the number of successful samples divided by total sample number. A sample prediction is regarded as successful if the DockQ of the best candidate structure is greater than 0.23.
b. We exclude 4 samples with surface antibody restraints, on which ClusPro fails to give a prediction.

**Figure 6.**
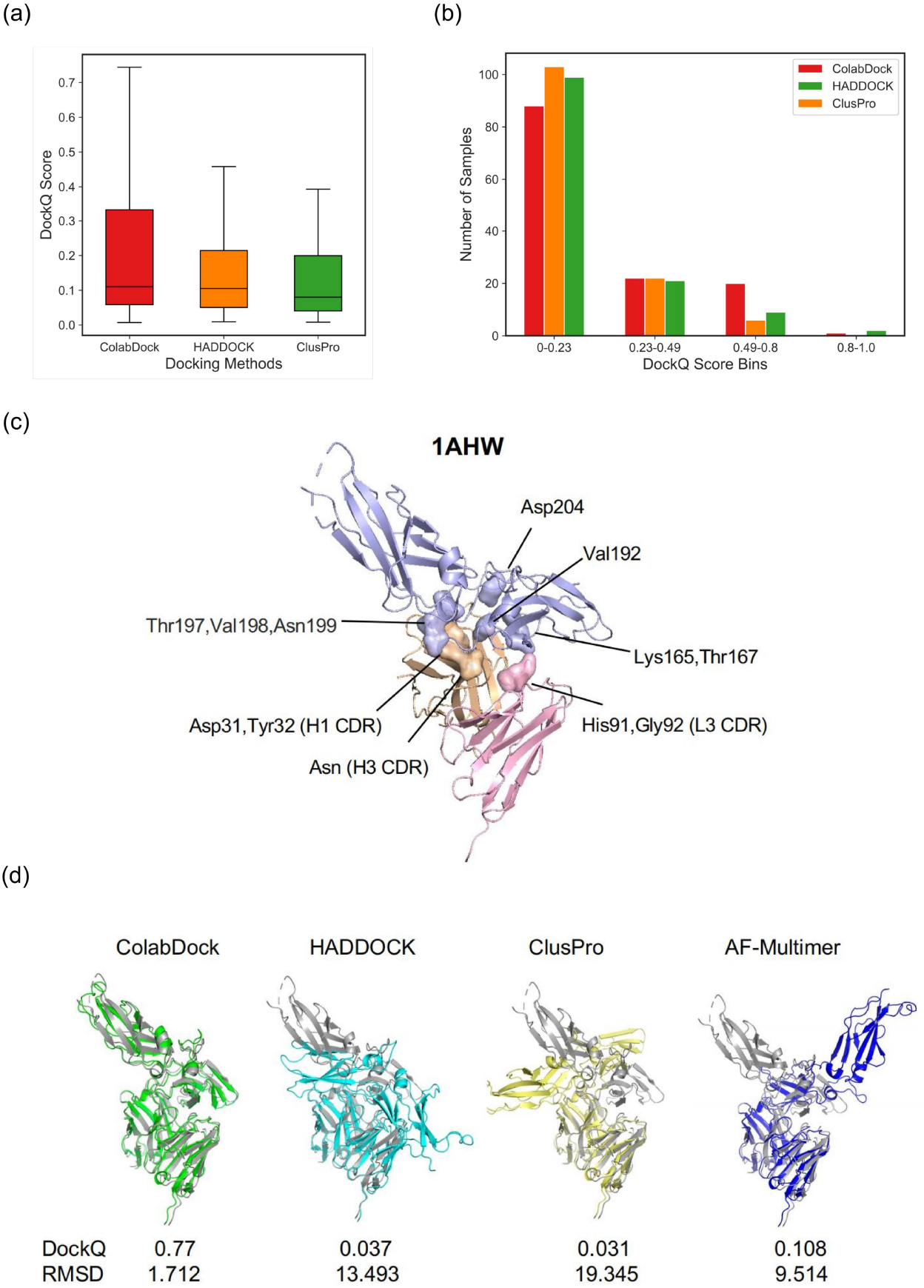
Comparison of ColabDock, HADDOCK and ClusPro on the simulated antibody-antigen benchmark set. **a**. DockQ of top1 structure predictions for the 131 samples by three docking methods with antibody-antigen surface restraints. ColabDock is better than the other two methods on the simulated antibody-antigen benchmark set **b**. Barplot counts the numbers of top1 structures predicted by three docking methods that fall into the four DockQ quality categories corresponding to below acceptable, acceptable, medium. and high quality. **c**. Interface of 1AHW (pdb code) and the sampled surface restraints. The tissue factor, VH, and VL are in light blue, wheat, and light pink, respectively. **d**. Top1 structures generated by ColabDock, HADDOCK, ClusPro, and AlphaFold-Multimer are aligned to the experimental structure for comparison. DockQs and RMSDs are listed for each predicted complex.

1AHW is a human TF(tissue factor)-antibody(5G9) complex, participating in the process of blood coagulation protease cascade. In this case, we randomly sample 5 interface residues (His91, Gly92 for the light chain, Asp31, Tyr32, Asn100 for the heavy chain) from the antibody and 7 interface residues (Lys165, Thr167, Val192, Thr197, Val198, Asn199, Asp204) from the TF (Figure 6c). These sampled residues in the antibody are mainly distributed on L1 CDR, H1 CDR, and H3 CDR loops. The sampled residues in TF are very close to important residues (Tyr156/157, Lys 165/166, Lys169, Arg200, and Lys201) revealed by the pervious mutation assay [21]. We use the three different methods integrated with above sampled restraints and AlphaFold-multimer to predict the interaction of TF and 5G9. As is shown in Figure 6d, ColabDock captures most of the native contacts at the interaction interface of 5G9 and TF, with DockQ of 0.77 and RMSD of 1.172, while the protein complex structures generated by the other methods largely differ from the native conformation. In contrast, top1 structure generated by HADDOCK tends to pull the Thr197, Val198, Asn 199 in 5G9 to bind the L1 CDR loops, while in the native structure, these three residues are in contact with H1 CDR loops. This structure satisfies 6 of 12 surface restraints and is evaluated as low-energy conformation by its score function. ClusPro only uses restraints in the structure filtering process instead of structure sampling. The top1 structure generated by ClusPro places Thr197, Val198, Asn 199 in 5G9 close to Asp31, Tyr32 (H1 CDR) and Asn100 (H3 CDR). In this case study, the score function of ColabDock better distinguishes between good and bad structures than that of the other two methods.

## Conclusion and Discussion

Protein docking provides crucial structure information to understand biological mechanisms. Despite the rapid development of deep models in protein structure prediction, most models make predictions in a free-docking way, which can potentially result in inconsistency between the experimental restraints and the predicted structure. To make better use of experimental results, in this study, we propose the ColabDock framework to incorporate the experimental restraints in protein complex structure modeling. The method is based on AF2 and generates complex structures in accord with the provided restraints through optimizing the input sequence. ColabDock is able to deal with various types of experimental restraints and offers an approach alternative to the traditional methods, which are based on FFT and energy functions.

In this work, we mainly make use of two types of restraints, the 1v1 and MvN restraints. The former is on the residue-residue level and corresponds to restraints such as those derived from XL-MS, while the latter is on the interface level, related to NMR and covalent labeling experiments. The results on the synthetic datasets show ColabDock achieves satisfying performance. Besides, as expected, with the increase of the number of restraints, the performance of ColabDock improves. Compared to HADDOCK and ClusPro, ColabDock achieves prominent performance when the quality of restraints is high. On two experimental datasets, ColabDock still outperforms HADDOCK and ClusPro regardless of the quantity and quality of the provided restraints. Lastly, we evaluated the performance of different docking methods on the antibody-antigen dataset. The proportion of medium or higher quality structures predicted by ColabDock is significantly higher than those of HADDOCK and ClusPro. This may show that ColabDock can refine the structures when it captures the rough relative orientation of protein complexes, and suggests the potential application of our model in antibody design.

Although ColabDock is based on AF2, it can be extended to any other fully differentiable protein structure prediction models, such as AlphaFold-Multimer and RoseTTAFold2. The reason we employ in this study AF2 is that AF2 was solely trained on protein domains and the protein complex structures were excluded. This choice guarantees the fairness in evaluation and comparison with the state-of-the-art docking models like HADDOCK and ClusPro.

Despite the success, ColabDock still has some limits. For now, ColabDock can only accept restraints on residue pairs at a distance below 22Å, which is determined by the upper limit of the distogram in AF2. This limitation makes the model only suitable for a fraction of the XL-MS reagents. ColabDock can also only handle complexes < 1200 residues on the NVIDIA A100 GPU due to memory consumption in optimization, and can be time-consuming especially for large protein complexes. The use of the bfloat16 floating-point format version AF2 is expected to help to save memory and accelerate the computation.

## Methods

### Datasets

In this work, we constructed a synthetic dataset and an experimental dataset to evaluate the performance on the simulated and experimental restraints, respectively.

#### Synthetic dataset

The synthetic dataset is curated from the protein docking benchmark 5.5 [22]. This benchmark consists of 271 protein complexes, with functions including enzyme-inhibitor, enzyme-substrate, antibody-antigen, and each has experimentally resolved structure. After removing structures with resolution > 3 Å, a synthetic dataset is constructed containing 241 protein structures. The synthetic dataset is further split into the benchmark set and the evaluation set. The former is used to compare the performance of ColabDock with HADDOCK and ClusPro. The latter is used to tune the hyperparameters and perform ablation studies. The details are provided as follows.

##### The benchmark set

The motivation of ColabDock is to incorporate experimental restraints into protein complex structure modeling to avoid inconsistency between experiment and prediction. Since a number of samples in protein docking benchmark 5.5 are in overlap with the AlphaFold-Multimer training set, we retrieved protein complexes with DockQ<0.23 for the prediction of AF2-multimer from the synthetic dataset, resulting in 84 complexes. We further removed two common (non antibody-antigen) protein complexes since ColabDock cannot process protein complexes with lengths more than 1200 residues at present. These proteins are composed of 37 common protein complexes and 45 antibody-antigen complexes, which are referred to as protein-protein benchmark and antibody-antigen benchmark, respectively. The performance of ColabDock is then evaluated on these ‘hard’ complex structures, along with HADDOCK and ClusPro.

##### The development set, the validation set, and the segment set

After the curation of the benchmark set, we first establish the development set by randomly sampling 30 proteins from the remaining 157 proteins. Then, from the remaining 127 proteins, 111 proteins with length below 700 are collected to compose the validation set. Meanwhile, we also retrieve proteins with length above 600 to build the segment set. The final segment set includes 29 proteins. It should be noticed that the validation set and the segment set have overlap.

#### Experimental datasets

We construct two datasets containing proteins with experimental restraints to evaluate the performance of the docking methods, i.e., the chemical shift perturbation (CSP) set and the covalent labeling (CL) set. CSP detects contact residues according to the change of chemical shifts upon binding to interacting proteins. CL labels the side chains of solvent accessible amino acids with reagents and differences in modification ratio between bound and unbound state can reveal residues in the interacting surface, which can be readout through mass spectrometry experiments.

Here, the CSP set consists of two proteins (1GGR and 3EZA) collected from Dominguez [5]. The CL set contains three proteins (1YAG, 2F8O, and 4INS) from Drake [19]. It should be noticed that protein 4INS has 3 biological assemblies, which consist of 4, 8, and 12 chains, respectively. Each biological assembly possesses a set of restraints. For the assembly with 4 chains, if chain A and B are regarded as a subunit, while chain C and D are regarded as another one, then the experimentally derived restraints are distributed between these two subunits. Thus, this assembly is denoted as 4INS_AB_CD. Similarly, the assemblies with 8 and 12 chains are denoted as 4INS_ABCD_EFGH and 4INS_ABCDEFGH_IJKL, respectively. For short, we call 4INS_AB_CD, 4INS_ABCD_EFGH and 4INS_ABCDEFGH_IJK as 4INS4, 4INS8, 4INS12 in this article, respectively.

### Restraints sampling

We classify restraints into two types according to the information they provide. The first is the 1v1 restraint in which the distance between a pair of residues are below a certain threshold, mimicking the cross linking & mass spectrometry data. Another type of restraint is referred to as the MvN restraint. It gives a series of restraints and each of them points out that a residue in one chain is in contact with a set of residues in another chain. It may occur that only part of the restraints is satisfied in the native structure. NMR Chemical Shift Perturbation (CSP), Paramagnetic Relaxation Enhancements (PRE), covalent labeling, and Deep mutation scanning (DMS) fall into the latter category. Additionally, considering the distances of part of the experimental restraints are above 8Å, we add loose restraints to both 1v1 restraints and MvN restraints. For antibody-antigen complexes, we adopt a sampling strategy to generate more dispersed restraints.

#### 1v1 restraints

1v1 restraints are sampled from clustered residue pairs in contact. Since restraints located close in the 3D space give similar structural information, we prefer to sample restraints that are dispersedly distributed on the interface. We first collect all the interchain residue pairs where the distances between the Cβ are below 8Å. Then, hierarchical clustering analysis is performed based on the residue index distance. If we denote the amino acid in the 12nd position in chain A as 12A, then the restraint between 12A and 15B is denoted as 12A-15B. The residue index distance between restraints 12A-15B and 22A-16B can be calculated as (22-12)+(16-15), equal to 11. Finally, restraints are randomly picked from as different clusters as possible. In ColabDock analysis, we sampled 2, 3, or 5 pairs of 1v1 restraints.

#### MvN restraints

The MvN restraints are derived from the 1v1 restraints. Suppose two 1v1 restraints 12A-15B and 14A-25B are given, we first expand each residue in the restraints by incorporating the 4 residues in the neighborhood. For example, 12A is expanded to residues from 10A to 14A. Then, for chain A, there are 7 residues (10A to 16A) derived from these 1v1 restraints, while for chain B, there are 10 residues (13B to 17B, and 23B to 27B). Finally, the derived MvN restraint means two of the 7 residues are in contact with at least one residue in the 10 residues.

#### Loose restraints

To evaluate performance of ColabDock on proteins containing restraints at longer distances, we additionally sample loose restraints. We first retrieve the interchain residue pairs with distances between 8Å and 20Å. Then, similar clustering, selection, and expansion processes are performed as in the generation of the 1v1 restraints. These restraints are referred to as loose restraints.

#### Antibody interface restraints

To mimic the DMS data of antibody-antigen complexes, for each chain, we first select residues that are in contact with the other chains. Then, 5-10 residues are randomly sampled from the selected residues in each chain to form the antibody interface restraints. These restraints are more dispersed than the MvN restraints.

### ColabDock

#### ColabDock model

AF2 is a deep neural network trained on experimentally solved and distilled protein structures. It takes as input the protein sequence, structure template, and multiple sequence alignment (MSA). The outputs of AF2 include the predicted atom coordinates, distogram, pLDDT, and pAE. Assume the length of the protein is *L*, the distogram is an *L*×*L*×64 matrix, where the (*i, j*, :) elements indicate the distance distribution between residue *i* and residue *j* in the 3D space. pLDDT and pAE are two quality measures with size of *L* and *L*×*L*, respectively. AF2 not only performs well on the protein structure prediction task, but has also been adopted for applications in protein design. ColabDesign [17], an AF2-based protein design protocol, has been applied in fixed-backbone design, protein hallucination [17], and peptide design [23].

Based on ColabDesign, we develop the ColabDock model, aiming to incorporate various experimentally derived restraints into the protein docking task. As is shown in Figure 1, ColabDock is composed of two stages, i.e., the generation stage and the prediction stage. In the generation stage, the model takes as input the templates of each chain in the complex, and generates a complex structure that is in accord with the provided restraints, through updating the input sequence. The index of the beginning residue is increased by 50 in each protein chain to minimize the impact of inter-chain positional encoding, as is adopted from the ColabFold [24] model. The input sequence is in the real number field rather than in the one-hot format. This has been proven to accelerate optimization convergence [17]. In the prediction stage, ColabDock predicts a complex structure based on the templates of each chain and the generated complex template. In both generation and prediction stages, the “aatype” variables in AF2 are set to the wild type sequence. This makes sure the sequences of the structures derived from both stages are in accord with the wild type. Two variables “target_feat” and “msa_feat” in the prediction stage are randomly selected from the wild type sequence and the optimized logit sequence from the generation stage with equal probabilities. The proposed ranking algorithm described below then sorts all the structure candidates. This setting achieves better performance according to our local experiments.

ColabDock adopts 4 types of losses in the generation stage, i.e., the monomer distogram loss, the restraint loss, pLDDT loss, and ipAE loss. The formats of these losses are as follows:

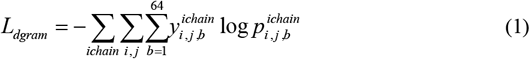

where *ichain* is the index of protein chains in the complex. *i, j*, and *b* are the indices of amino acids and bins in the distogram. 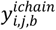 and 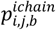 represent the one-hot encoded distance and the predicted distance probability, respectively. The restraint loss has similar format to the monomer distogram loss:

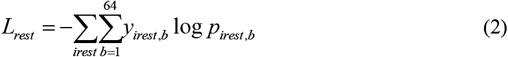

For 1v1 restraints, the loss is calculated directly. For MvN restraints, we first sort along the N dimension and select the most probable interacting residue for each of the M residues. Then, we sort along the M dimension and select a number of interacting residue pairs to optimize. *pLDDT* and *ipAE* losses are defined as:

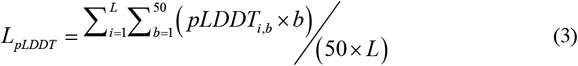

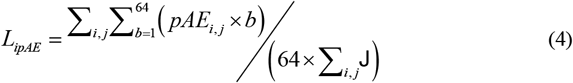

where 𝕁 is a matrix in which the entity is set to 0 if *i* and *j* belong to the same chain, and otherwise set to 1. The intra-chain distogram loss guarantees the predicted chain conformations are in accord with the input templates. The restraint loss brings the residues in the experimental restraints close. pLDDT and ipAE make sure the predicted complex structure is of high quality. The final loss is the weighted sum of these 4 types of losses:

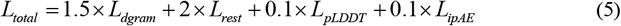

During optimization, the distance of the provided restraints is by default set to 8Å. This value can be adjusted according to the types of the restraints. The learning rate is set to 0.1.

#### Segment based optimization

Since ColabDock makes predictions through backpropagation, it will consume large GPU memory and limit the feasible protein length. To handle longer proteins, we propose the segment based optimization protocol. Specifically, a set of residues is first cropped out from the whole sequence before each optimization step starts. With 50% probability, the cropped residues contain provided restraints and the optimization loss includes all the 4 items. For the remaining 50% probability, the cropped residues are randomly chosen and the loss only contains *L*_*dgram*_, *L*_*pLDDT*_, and *L*_*ipAE*_. During the optimization, only the representation of the cropped residue is updated. The remaining part stays unchanged. Our experiments show that for most proteins, if the number of cropped residues is set to 200, then 100 optimization steps are enough to reach a convergent structure (Figure S6). We use this setting as the default setting.

#### Ranking algorithm

It is observed that the performance of ColabDock is largely influenced by the stochastics in initialization and optimization. To alleviate this, for each protein, ColabDock is run multiple times to generate diverse docking structures. Hereafter, we refer to the number of runs as **round**. In each round, the number of backpropagations is referred to as **step**. The generated structures from all the rounds are sorted according to the proposed ranking algorithm.

The ranking algorithm is developed based on the RankingSVM (RSVM) model [25]. It takes 5 types of features as input, including ipTM, contact number, contact pLDDT, number of satisfied restraints, and average error. ipTM is the output of AF2, measuring the quality of the interface in the complex. Contact number is defined as the number of interchain residue pairs in the predicted structure whose distances are below 8Å. Contact pLDDT is the average pLDDT of the residues in contact. The number of satisfied restraints indicate whether the predicted complex structure is in accord with the provided restraints. Average error only considers the residue pairs in the provided restraints. The error is set to the excess if the distance of the residue pair is larger than 8Å, otherwise it is equal to 0.

We establish two RSVM models to sort the structures generated in each round and those selected from all the rounds, which are referred to as the intra-RSVM and the inter-RSVM, respectively. The RSVM models are trained on the development set. Training details are described in support information. In practical application, intra-RSVM is first used to select the top 5 structures in each round. Then, inter-RSVM is applied to rank all the selected structures.

### Evaluation measures

To evaluate the performance, we mainly adopt two measures, i.e., DockQ [18] and Cα-RMSD. The former focuses on the quality of the interface, while the latter measures the global structural difference. Structures can be divided into 4 classes according to DockQ: incorrect (0<DockQ<0.23), acceptable (0.23<= DockQ<0.49), medium (0.49<=DockQ<0.80), and high (DockQ>0.80). We regard complex structures with DockQ>=0.23 as correct predictions. DockQ is calculated using the official code (https://github.com/bjornwallner/DockQ).

## Acknowledgement

We thank George Jones from Vajda lab for very helpful discussions on usage of ClusPro.

## Data and Code Availability

The ColabDock code will be available at https://github.com/JeffSHF/ColabDock.

